# Amyloid beta peptides (Aβ) from Alzheimer’s disease neuronal secretome induce endothelial activation in a human cerebral microvessel model

**DOI:** 10.1101/2022.07.27.501634

**Authors:** Yu Jung Shin, Kira M. Evitts, Solhee Jin, Caitlin Howard, Margaret Sharp-Milgrom, Jessica E. Young, Ying Zheng

**Affiliations:** Department of Bioengineering; University of Washington; Seattle, WA, 98109; United States of America; Institute for Stem Cell and Regenerative Medicine, University of Washington, Seattle, WA 98109, United States of America; Department of Laboratory Medicine and Pathology, University of Washington, Seattle, WA 98109, United States of America

**Keywords:** Alzheimer’s disease, amyloid beta peptide (Aβ), endothelial activation, vascular dysfunction, disease modeling, 3D microvessels

## Abstract

In Alzheimer’s disease (AD), secretion and deposition of amyloid beta peptides (Aβ) have been associated with blood-brain barrier dysfunction. However, the role of Aβ in endothelial cell (EC) dysfunction remains elusive. Here we investigated AD mediated EC activation by studying the effect of Aβ secreted from human induced pluripotent stem cell-derived cortical neurons (hiPSC-CN) harboring a familial AD mutation (Swe^+/+^) on human brain microvascular endothelial cells (HBMECs) in 2D and 3D perfusable microvessels. We demonstrated that increased Aβ levels in Swe^+/+^ conditioned media (CM) led to stress fiber formation and upregulation of genes associated with endothelial inflammation and immune-adhesion. Perfusion of Aβ-rich Swe^+/+^ CM induced acute formation of von Willebrand factor (VWF) fibers in the vessel lumen, which was attenuated by reducing Aβ levels in CM. Our findings suggest that Aβ can trigger rapid inflammatory and thrombogenic responses within cerebral microvessels, which may exacerbate AD pathology.

## 1. Introduction

Alzheimer’s disease (AD) is a progressive neurodegenerative disease and the most common form of dementia (*1*, *2*). The neuropathological hallmarks of AD include the deposition of extracellular amyloid beta (Aβ) plaques, primarily of the Aβ 1-42 isoform (*3*), formation of intracellular neurofibrillary tangles (NFTs), and progressive loss of neurons and synapses. Decades of research have been devoted to understanding the mechanisms of AD development and potential treatments, but only a few FDA-approved agents for the treatment of AD are available and no disease-modifying treatments are available (*4*). Existing AD research has primarily focused on dysfunctions of cell types in the central nervous system (CNS) such as neurons, astrocytes, and microglia, but growing evidence suggests a role for vascular dysfunction in the development and progression of AD (*4*). Greater than 90% of patients with Alzheimer’s disease (AD) have been found with amyloid deposits along their cerebral vasculature, a condition known as cerebral amyloid angiopathy (CAA) (*5*). Aberrant vascular morphology (increased tortuosity of vessels, increased presence of string vessels, thinning of vessel walls, etc.), degeneration of vessels in affected brain regions, and decreased cerebral blood flow (CBF) are also apparent in AD (*6*). However, the driving factors for vascular malformations present in AD pathogenesis remain unclear.

*In vivo*, brain microvessels form a vast and complex hierarchical network to regulate transport of nutrients and prevent toxic pathogens and metabolites from entering the CNS. As the interface between circulation and brain tissue, endothelial cells (ECs) play a critical role in building a strong blood-brain barrier (BBB). In their healthy state, brain ECs express tight junction proteins to limit the diffusion of molecules into the brain (*7*); when activated in response to pro-inflammatory signals, brain ECs exhibit (1) loss of vascular integrity, (2) increased expression of adhesion molecules, such as intercellular adhesion molecule 1 (ICAM-1) and vascular cell adhesion molecule 1 (VCAM-1), (3) upregulation of human leukocyte antigen molecules, (4) exocytosis of Weibel-Palade bodies (WPB) containing P-selectins and von Willebrand Factor (VWF), indicating a prothrombotic state following endothelial injury, (5) increased secretion of cytokines (IL-8, IL- 12), and (6) assembly of stress fibers (*8*–*10*).

In patients with AD, serum and plasma analysis show increases in Aβ proteins in the systemic circulation, along with plasma biomarkers of EC activation such as VWF (*6*, *11*). Increases in plasma levels of soluble adhesion proteins such as E-selectin, P-selectin, ICAM-1 and VCAM-1 have also been found in AD patients relative to controls (*12*–*15*). Nevertheless, studies of human AD-mediated vascular dysfunction are limited as it is challenging to study cellular function non-invasively in the human brain. *In vitro* cultures of EC monolayers have shown that exogenously applied Aβ peptides led to nuclear and mitochondrial DNA damage, cell death, increased BBB permeability and increased monocyte adhesion on ECs (*4*, *8*, *16*, *17*). In bovine aortic EC cultures, Aβ 1-42 was found to induce endothelial dysfunction by raising intracellular calcium levels via calcium-permeable Aβ channels (*18*). These 2D *in vitro* studies, however, lack proper vascular architecture, lack *in vivo* flow characteristics, and apply exogenous Aβ at micromolar doses, much higher than the Aβ oligomer levels in the human brain measured at picomolar concentrations (*19*). It remains elusive to what extent Aβ peptides drive human brain microvascular injuries and dysfunctions in AD conditions.

Transgenic mouse models of AD are often utilized to show cerebrovascular phenotypes, however, not all of the models show the same aberrations and they do not always occur at the same time (*20*). For example, decreased vessel density, a common vascular alteration in human AD, is observed in only one of the ten most common AD mouse models, and depending on the mouse line, the onset of vascular amyloid deposition can occur anywhere between 3 months and 24 months of age (*20*). Furthermore, these animal models often have non-physiologic expression of a number of AD-related transgenes. Hence there is a critical need to evaluate the role of Aβ in AD-associated vascular dysfunction in a system that structurally and functionally mimics brain vasculature and includes accurate Aβ levels.

More recently, advances in microfluidic-based techniques enabled modeling of the 3D BBB with perfusable lumens that can recapitulate the biochemical and biophysical perturbations occurring in cerebral blood vessels (*21*, *22*). Culturing ECs under flow improves EC homogeneity, homeostatic control, EC alignment with flow, vascular development, and EC proliferation (*23*–*25*). In primary HBMECs, flow has also been shown to increase the tightness of junctions in vascular models, which significantly decreased EC permeability, better reflecting the highly impermeable barrier of the *in vivo* BBB (*26*). Hence, 3D perfusable human cerebral vascular models provide timely opportunities for studying the responses of human brain ECs and microvessels to the increased levels of secreted Aβ in AD in physiological and pathological-relevant conditions.

In this study, we utilized primary HBMECs and established a 3D perfusable human brain microvessel platform to study AD-associated endothelial activation. We modeled the AD response by perfusing these microvessels with neuronal secretomes, collected as conditioned medium (CM) from human neuronal cultures. These neurons were differentiated from human induced pluripotent stem cells (hiPSCs), that harbor the amyloid precursor protein (APP) Swedish mutation KM670/671NL (APPSwe). This mutation increases Aβ levels by enhancing amyloidogenic APP cleavage by β-secretase 1 (BACE1) and is causative for early-onset familial AD (FAD) (*27*, *28*). Cells with AD mutations, including the Swedish mutation, have been used to study blood-brain barrier (BBB) changes in AD and CAA (*29*, *30*). One recent study demonstrated that conditioned media (CM) collected from hiPSC-derived APPSwe neurons led to increased permeability in 2D monolayers of proliferating brain ECs, while WT CM did not affect permeability (*31*), suggesting that this CM study can recapitulate features of BBB dysfunction seen in AD brain.

We compared the endothelial alterations induced by treatment with CM from cortical neuron cultures differentiated from APPSwe hiPSCs (hiPSC-CNs) and their isogenic wild-type (WT) control cell lines. In 2D, treatment of HBMECs with CM from AD hiPSC-CNs led to cytoskeletal reorganization and upregulation of genes involved in immune cell interaction with ECs. In 3D flow-directed brain microvessels, perfusion of CM from APPSwe hiPSC-CNs elicited acute endothelial activation and formation of transluminal VWF fibers, suggesting the initiation of thrombotic microangiopathy within the luminal space. By showing reduced endothelial activation to CM from neurons deficient in APP expression (APP^KO^), or from APPSwe cells treated with a β-secretase inhibitor (BACEi), we demonstrated the pivotal role of Aβ in EC activation. Our findings highlight that increased Aβ levels secreted by AD neurons lead to endothelial activation in brain microvessels. This suggests that Aβ has a direct role in eliciting acute hemostasis and vascular inflammation in the cerebrovasculature of AD patients.

## 2. Results

### 2.1 APPSwe hiPSCs-CNs secretes Aβ in a genotype-dependent manner

To investigate the effect of AD neuron derived paracrine signaling on HBMECs, we differentiated hiPSCs from an established cell line harboring APPSwe mutation into cortical neurons (CNs) (*28*, *29*) (Figure 1A,B). Following a previously published protocol (*32*, *33*), neural progenitor cells (NPCs) were first differentiated from hiPSCs for 3 weeks, followed by another 3 weeks of differentiation into neurons. At the end of the 3-week differentiation, CNs were enriched in the cultures, as marked by robust immunofluorescent staining for the neuron marker MAP2 (Figure 1B). hiPSC lines homozygous (Swe^+/+^) and heterozygous (Swe^+/WT^) for the APPSwe mutation were used to generate hiPSC-CNs (Figure 1A). Control hiPSC-CNs were made using the parental wild type (Swe^WT/WT^) line for the APPSwe lines, and an APP knock out (APP^KO^) cell line was used as an additional control (*28*, *34*).

**Figure 1.**
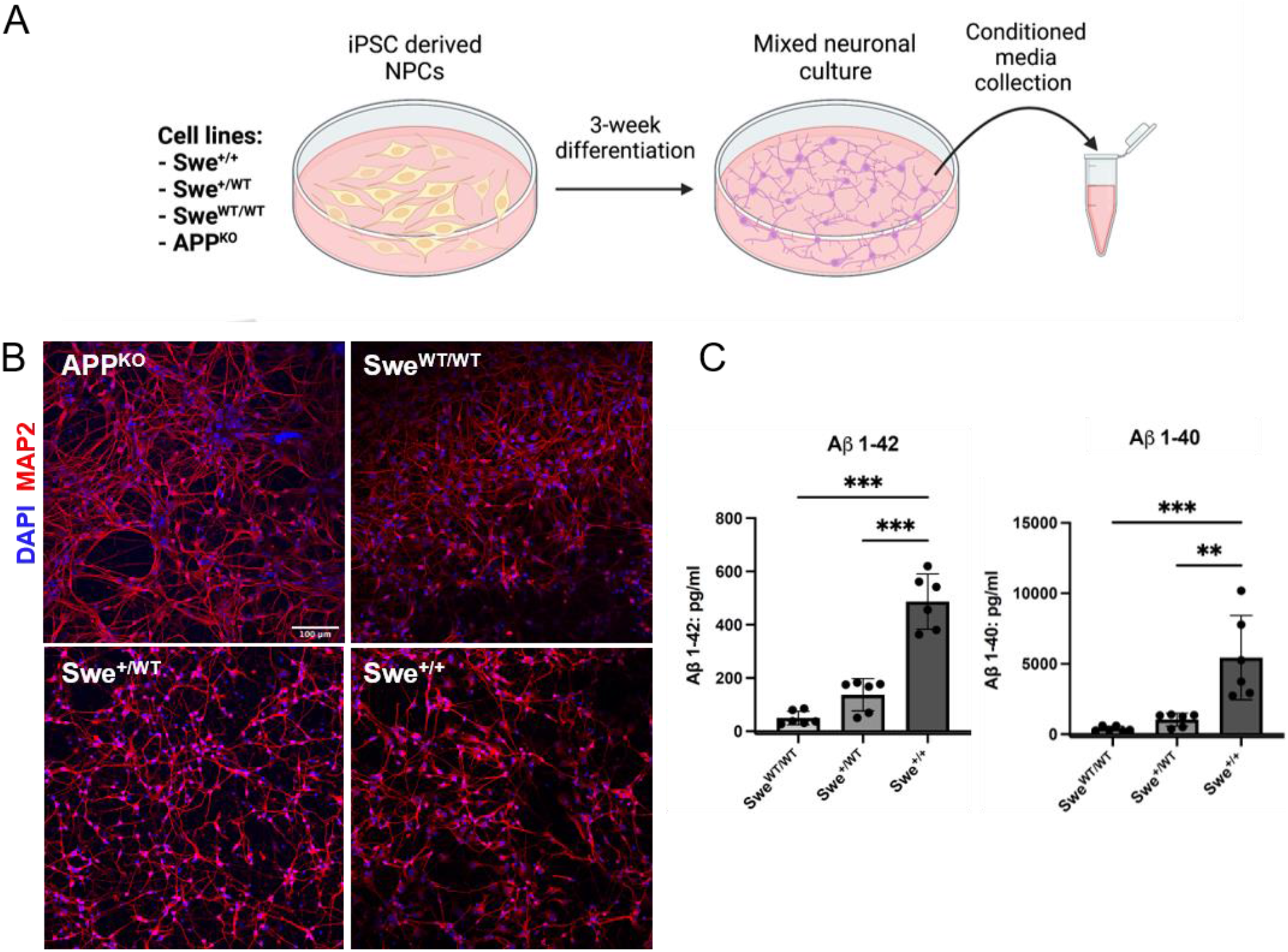
Condition media (CM) collection and characterization of hiPSC-CN cultures. **(A)** Schematic overview of CM obtained from neuronal cultures. Swe^+/+^ (homozygous) and Swe^+/WT^ (heterozygous) CNs were differentiated from hiPSCs harboring the APP Swedish mutation. For our controls, CNs were differentiated from an isogenic control line, Swe^WT/WT^, and a line that has CRISPR knockout of the APP gene, APP^KO^. **(B)** Immunofluorescent (IF) images of cortical neurons differentiated from APP^KO^, Swe^WT/WT^, Swe^+/WT^, and Swe^+/+^ hiPSCs show enriched staining for cortical neurons (MAP2, red). Scale bar: 100 μm **(C)** ELISA quantification of CM shows increased levels of Aβ expression in APPSwe CNs. n=3 biological replicates. Error bars, mean ± SEM. *p < 0.033, **p < 0.002, and ***p < 0.001 by one-way ANOVA with Tukey’s correction for multiple comparisons test.

Following 3-weeks of differentiation, conditioned media (CM) was collected from hiPSC-CN cultures and examined for their Aβ 1-42 and 1-40 levels using an ELISA assay (Figure 1C). CM collected from Swe^+/+^ and Swe^+/WT^ hiPSC-CNs showed increased Aβ 1-42 and 1-40 levels compared to controls, with the highest mean in the Swe^+/+^ line for Aβ 1-42 (487 pg/mL) and 1-40 levels (5431 pg/mL). The Swe^+/WT^ line showed significantly lower Aβ 1-42 (137 pg/mL) and Aβ 1-40 (1033 pg/mL) levels compared to Swe^+/+^, but significantly higher than that collected from the Swe^WT/WT^ (Aβ 1-42 at 50 pg/mL and 1-40 at 385 pg/mL). These results confirm that the APPSwe neurons secrete increased levels of Aβ and may be a viable way to mimic paracrine signals from neurons in the AD brain.

### 2.2 Paracrine signaling from neurons induces endothelial activation and cytoskeletal reorganization in 2D monolayer culture

Under stress, endothelial cells undergo EC activation, which leads to the formation of stress fibers (*10*). We next treated HBMECs in a 2D monolayer (Figure 2A (i)) with CM obtained from hiPSC- CN cultures to evaluate the paracrine effect of neurons on HBMECs. Four media groups were tested to assess endothelial activation after a 6-hour treatment: Swe^+/+^ CM, Swe^+/WT^ CM, Swe^WT/WT^ CM, and unconditioned media (UCM). Upon treatment with Swe^WT/WT^ CM, HBMECs exhibited quiescent cytoskeletal structures characterized by a cortical actin rim around the peripheral junctions (Figure 2A iii-arrows). In comparison, Swe^+/+^ CM and Swe^+/WT^ CM treated HBMECs show reorientation of actin fibers into elongated stress fibers with increased cell polarity (Figure 2A iv and v-arrowheads), suggesting their activated state. In particular, Swe^+/+^ CM treated HBMECs show significant retraction of endothelial cells (Figure 2A (v-arrowheads)), resulting in loss of junctions between cells, and increasing gap formation between adjacent endothelial cells.

**Figure 2.**
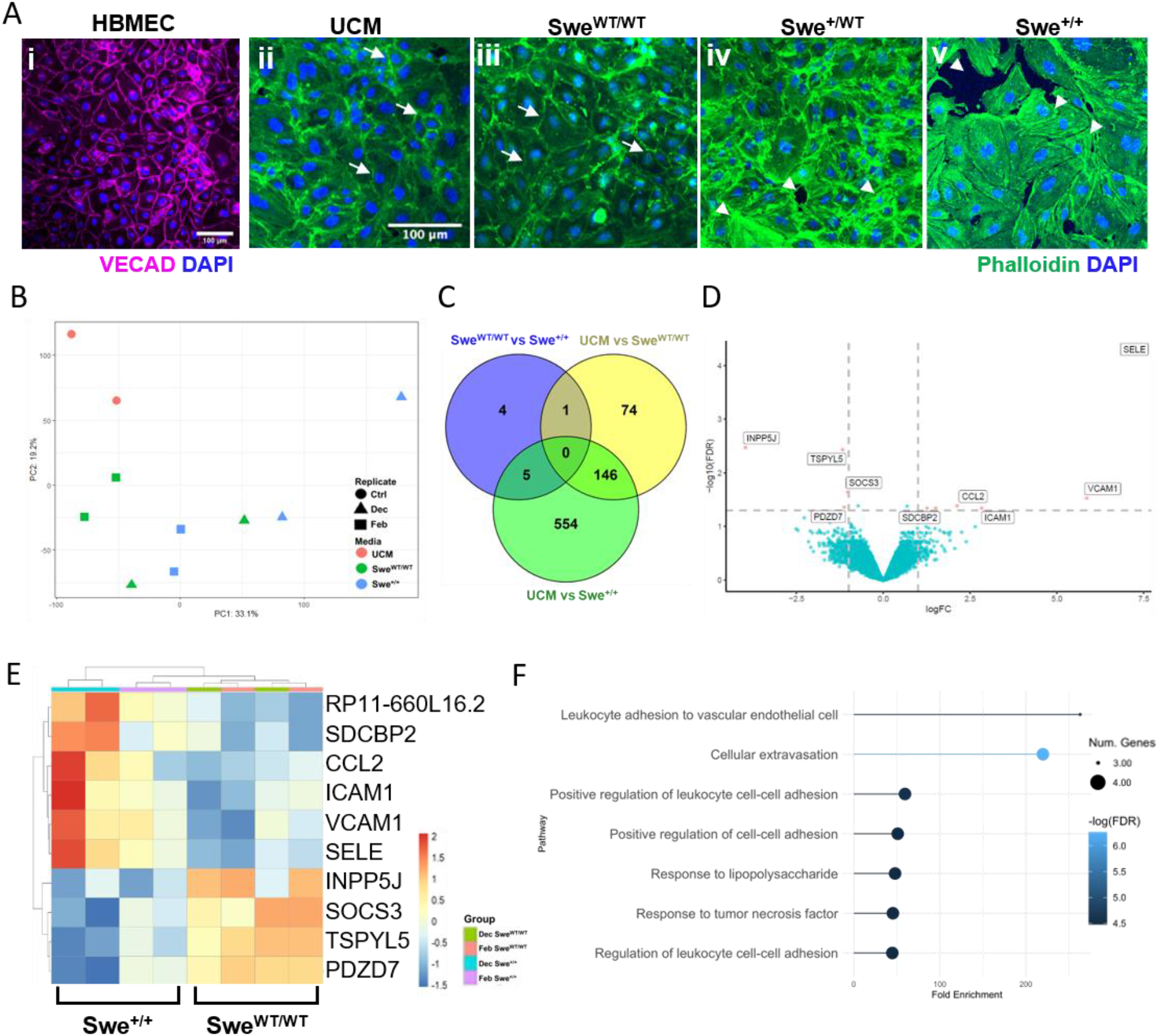
Cytoskeletal and transcriptomic changes in 2D HBMEC cultures after CM treatment. **(A)** Representative IF images of 2D cultured **(i)** untreated HBMECs and **(ii-v)** HBMECs treated with CM media. 2D HBMEC IF staining shows good vascular endothelial cadherin junction (VECAD, magenta) under normal culture conditions. **(ii)** 2D cultures exposed to Swe^WT/WT^ media maintain cortical actin cytoskeletal staining (Phalloidin, green) as denoted by white arrows **(iii)**, Swe^+/WT^ CM treatment shows endothelial retraction and f-actin stress fiber formation (Phalloidin, green) **(iv)** which is intensified in Swe^+/+^ treatment **(v)** as denoted by white arrow heads. Representative images from n=3 biological replicates. Scale bar: 100 μm **(B)** PCA of RNA-seq data from HBMECs treated with three media types (UCM, Swe^WT/WT^ and Swe^+/+^) in 2D. **(C)** Venn diagram showing the overlap of genes significantly changed by CM media treatment. 10 genes were differentially expressed between Swe^WT/WT^ and Swe^+/+^ CM treatment. **(D)** Volcano plot showing DEGs between, Swe^WT/WT^ treated media and Swe^+/+^ treated media. **(E)** Heatmap of log counts per million (CPM) values for genes that were identified to be significant in Swe^+/+^ treated HBMECs. **(F)** Gene Ontology terminology analysis for biological processes involved in upregulated genes in Swe^+/+^ CM treated HBMECs.

### 2.3 RNA sequencing (RNAseq) reveals endothelial activation signatures in CM treated HBMECs

To understand the transcriptomic changes in endothelial cells following CM treatment, we sequenced RNA collected from 2D HBMECs that were treated with two batches of Swe^+/+^ CM, Swe^WT/WT^ CM and UCM. Principal component analysis (PCA) showed that individual UCM, Swe^WT/WT^ and Swe^+/+^ CM treated groups clustered together (Figure 2B). In particular, the UCM cluster is distinctly separated from the Swe^WT/WT^ and Swe^+/+^ clusters, suggesting that neuronal secretomes had a large effect on EC function compared to UCM. Clusters also form within the same media batch groups, with distinct cluster presentations for Swe^WT/WT^ and Swe^+/+^ groups. Batch-to-batch variation appeared to exist in the CM obtained, leading to overlaps among clusters, which is likely due to the inherent heterogeneity of hiPSC-CNs and secretome production (Supplemental Figure 1). For analysis of RNAseq, we combined both batches for unbiased representation of transcriptomes that were differentially expressed between Swe^WT/WT^ and Swe^+/+^ treated ECs. Results showed 10 genes that were differentially expressed between HBMECs treated with Swe^WT/WT^ CM and Swe^+/+^ CM (Figure 2C). This includes upregulation of genes associated with endothelial adhesion molecules: SELE (~7-fold change), VCAM-1 (~6-fold change) and ICAM-1 (~3-fold change) (Figure 2D and 2E). CCL2 and SDCBP2 were also upregulated, which supports the presence of higher levels of inflammatory cytokines in Swe^+/+^ CM. Down-regulated genes include: INPP5J, TSPYL5, SOCS3 and PDZD7 (Figure 2C and 2D). Downregulation of the TSPYL5 gene, known to be associated with endothelial proliferation and tube formation, suggests angiogenic ability may be attenuated in ECs with AD exposure (*35*). GO term and KEGG pathway analysis showed up-regulation of pathways related to leukocyte adhesion, cellular extravasation, and endothelial inflammation (Figure 2F). 24-hour treatment of HBMECs with Swe^+/+^ CM and Swe^WT/WT^ CM confirmed increased VCAM-1 expression on the endothelial surface upon stimulation with Swe^+/+^ CM (Supplemental figure 2).

Overall, both phenotypic and transcriptomic analyses of HBMECs suggest that ECs are activated by neuronal secretomes in AD CM through cytoskeletal reorganization, and transcriptional alteration. These data indicate that paracrine signaling from AD neurons elicits a pro-inflammatory, pro-thrombotic, and anti-angiogenic environment in brain ECs.

### 2.4 Swe^+/+^ conditioned media induces endothelial activation in 3D flow-mediated engineered cerebral microvessels

The endothelium *in vivo* forms an intact lumen and 3D network structure, responding to flow and cytokines synchronically. When activated, the endothelium is known to release granules, such as VWF, which can extend and multimerize into large fibers, under flow, to bind platelets and other blood cells (*21*). Such activation denotes a shift of the endothelium toward a prothrombotic phenotype and has been shown as a signature phenomenon in 3D microvessels (*21*). We examined whether Swe^+/+^ CM would induce rapid activation of 3D flow-mediated engineered cerebral microvessels. Using the engineered microvessel method previously developed by our lab (*36*), we created devices with a double grid geometry (8×8) for microvessels with a feature height of 150 μm (Figure 3A), and seeded with HBMECs at a concentration of 7×10^6^ cells/mL. After culture for 5-7 days under gravity-driven flow, a continuous and confluent endothelium was established in 3D engineered cerebral microvessels (Figure 3A).

**Figure 3.**
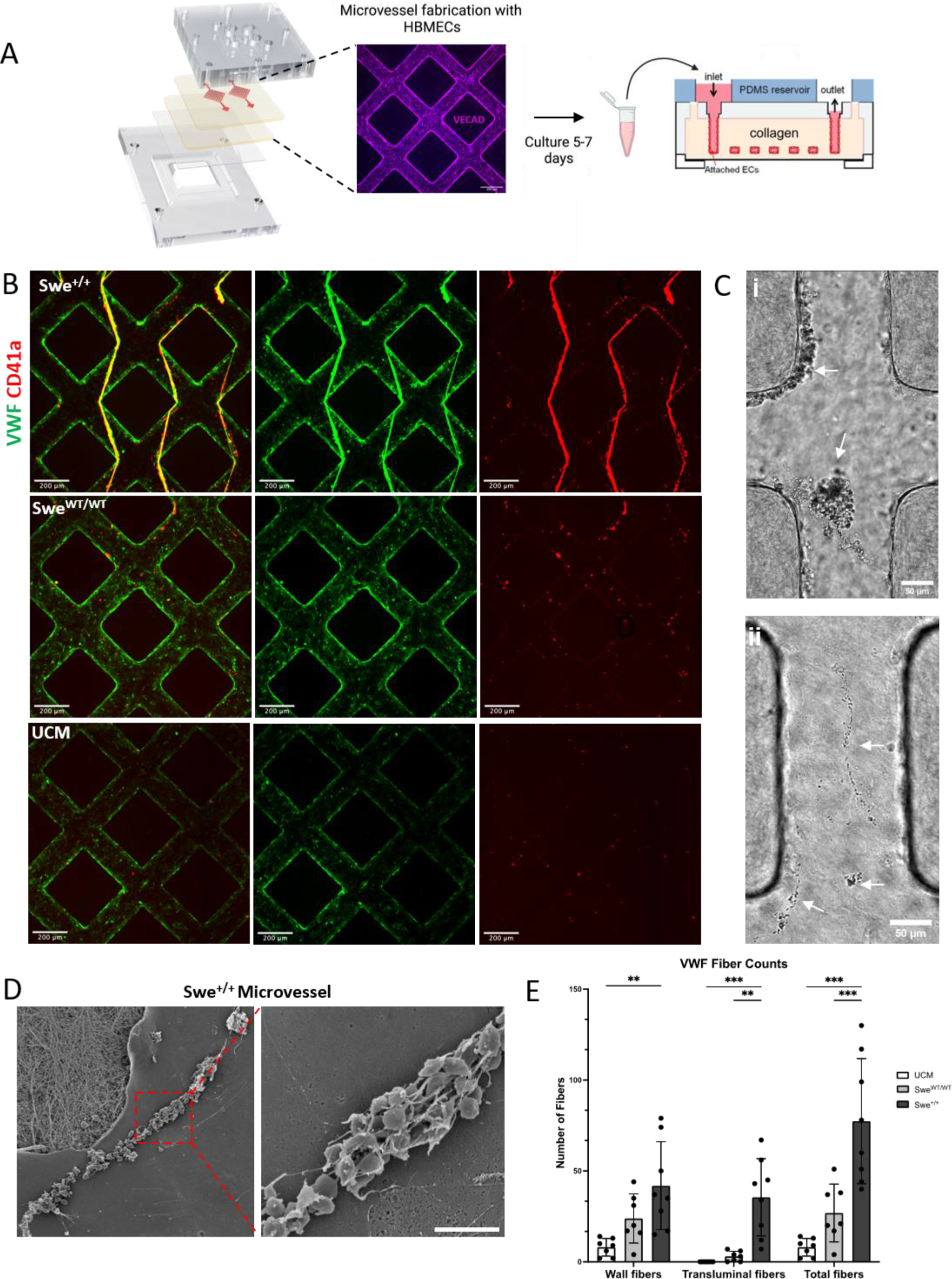
Perfusion of CM stimulates VWF release in 3D engineered cerebral microvessels. **(A)** Schematic illustrating engineered microvessel fabrication and treatment of vessels with CM. **(B)** Representative images of microvessels treated with CM (bottom: unconditioned base media (UCM), middle: Swe^WT/WT^, and top: Swe^+/+^). Swe^+/+^ CM perfusion causes VWF fiber formation (VWF, green) and induces binding of washed isolated platelets (CD41a, red). By comparison little to no VWF fiber formation or platelet binding occurs in UCM and Swe^WT/WT^ CM treated microvessels. Scale bar: 200 μm. **(C)** Brightfield images of platelets binding to VWF fibers in engineered cerebral microvessels. Platelets bound to both **(i)** transluminal fibers and **(ii)** wall fibers. Arrows indicate platelet binding and aggregation. Scale bar: 50 μm. **(D)** SEM image of microvessels containing a wall VWF fiber with bound platelets. Platelets are bound in a mesh of VWF fibers present on the luminal surface of microvessels. Scale bar: 5 μm. **(E)** Quantification of VWF fibers formed upon perfusion of engineered cerebral microvessels with CM. n=4 biological replicates. Error bars, mean ± SEM. *p < 0.05, **p < 0.01, and ***p < 0.001 by two-way ANOVA with Tukey’s correction for multiple comparisons test.

We then perfused cerebral microvessels with different CM for 1 hour under gravity-driven flow (Figure 3A) and evaluated the formation of VWF fibers in the vessel lumen. Treatment of vessels with Swe^+/+^ media led to the formation of large transluminal VWF fibers throughout the vessels (Figure 3B (green) and 3C), which was not observed upon perfusion with CM from the Swe^WT/WT^ line or UCM. The extended VWF fibers in the vessel lumen expose binding sites for platelets, which could initiate thrombosis in vessels (*37*). We next perfused washed platelets isolated from whole blood, stained for the platelet marker CD41a, through the microvessels. Upon perfusion, platelets readily bound to VWF fibers in the Swe^+/+^ condition, but there was minimal binding in vessels with no VWF fibers (UCM and Swe^WT/WT^) (Figure 3B (red)). Using VWF staining and platelet perfusion, two types of VWF fibers were identified in perfused microvessels, including transluminal fibers, seen as large fibers in the center of the vessel lumen, and wall fibers, as small fibers found on the wall of the vessels that are marked by platelet strings (Supplemental Figure 3). Brightfield and scanning electron microscopy of the Swe^+/+^ CM treated microvessels showed that large aggregates of platelets are bound to ultra-large transluminal VWF fibers (Figure 3C(i)) whereas single platelet binding is bound on wall string fibers. (Figure 3C(ii) and 3D). Quantification of wall and transluminal fibers in vessels perfused with Swe^+/+^ CM showed a significant increase in transluminal and total fibers, and a trend toward increased wall fiber formation across several replicates compared to controls (UCM and Swe^WT/WT^). (Figure 3E)

Furthermore, we observed that in brain microvessels perfused with Swe^+/+^ CM, leukocytes were able to bind to the endothelial surface and initiate a tethering and rolling response. This suggests that endothelial cells were activated by CM with higher Aβ concentrations which facilitated immunoadhesion within the microvessels (Supplemental Figure 4; Supplemental video 1). These results suggest that perfusion of engineered microvessels with CM with high Aβ levels can induce EC activation and VWF fiber formation that is not present in vessels perfused with CM from cell lines with low Aβ production.

### 2.5 Knocking out APP and inhibiting the beta-secretase enzyme BACE1 with a small molecule inhibitor (BACEi) attenuate endothelial cell activation and VWF release

To test whether the increased level of Aβ in our APPSwe CM is a driving factor for endothelial activation, we perfused microvessels with CM from Swe^+/+^, Swe^+/WT^, and Swe^WT/WT^ hiPSC-CNs treated with a BACE1 inhibitor (BACEi). BACEi inhibits BACE1 function and prevents the amyloidogenic cleavage of APP into Aβ (Figure 4A) (*38*, *39*). CM was collected from cultures of hiPSC-CNs after treatment with either BACEi (25 nM) or vehicle control (DMSO) for 72 hours (Figure 4B). ELISA measurements showed a statistically significant 44% decrease in Aβ 1-42 levels (DMSO: 170 pg/mL, BACEi: 75.4 pg/mL) and 45% decrease in Aβ 1-40 levels (DMSO: 1504 pg/mL, BACEi: 683 pg/mL) in Swe^+/+^ BACEi treated cultures compared to DMSO controls (Figure 4C and Supplemental Figure 5). In Swe^+/WT^ cultures, BACEi treatment led to a statistically significant 38% decrease in both Aβ 1-42 (DMSO: 101 pg/mL, BACEi: 38.1 pg/mL) and 1-40 (DMSO: 794 pg/mL, BACEi: 298 pg/mL) levels compared to controls. In Swe^WT/WT^ cultures, BACEi treatment led to a 28% reduction in Aβ 1-42 levels (DMSO: 32.4 pg/mL, BACEi: 9.14 pg/mL) and 26% reduction in Aβ 1-40 levels (DMSO: 264 pg/mL, BACEi: 69.1 pg/mL).

**Figure 4.**
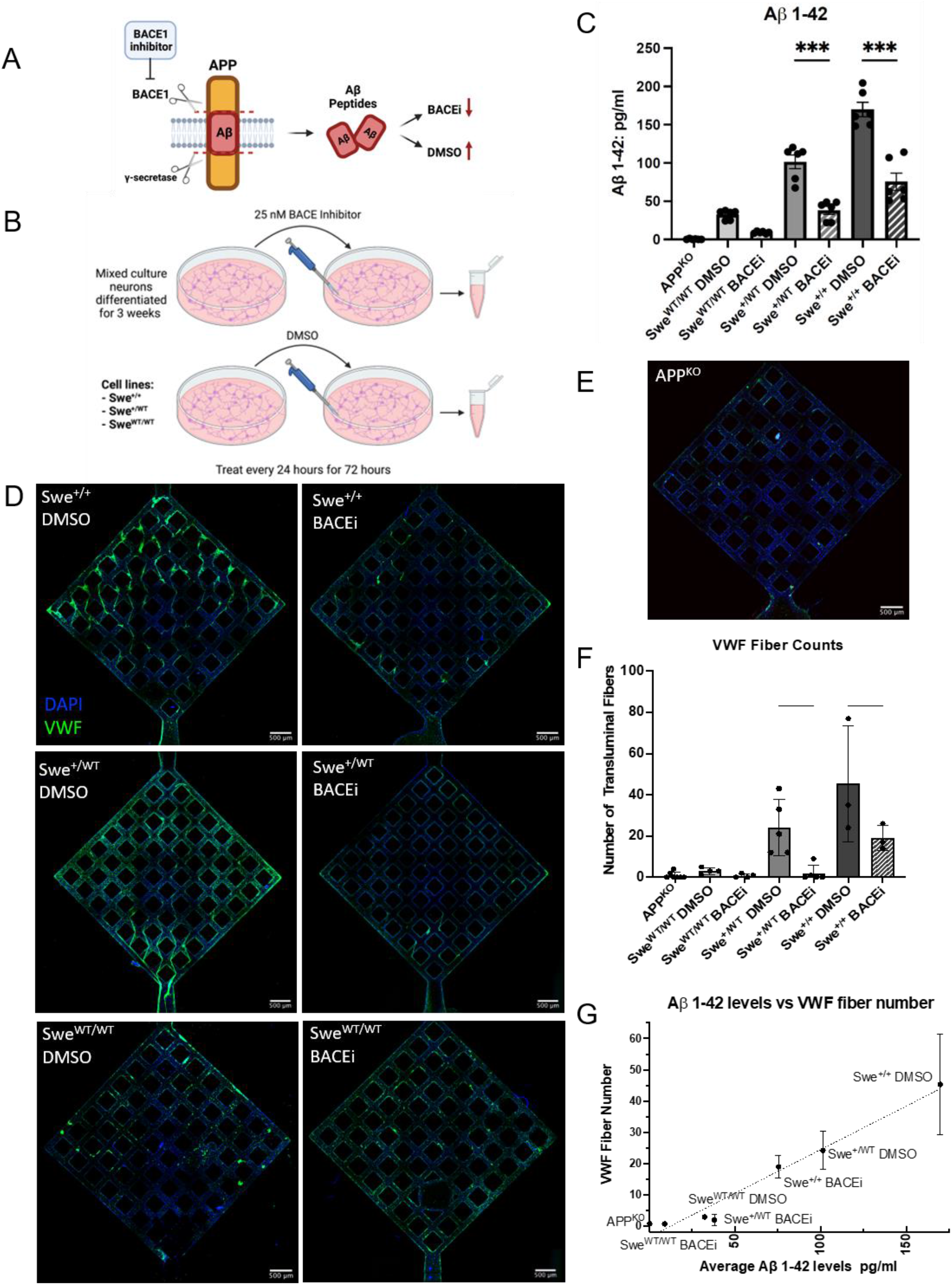
CM collected from BACE1 inhibitor (BACEi) treated neurons or APP^KO^ neurons cause reduced VWF fiber formation in 3D microvessels. **(A)** BACEi mechanism of action. BACEi inhibits BACE1 cleavage of APP and results in decreased production amyloidogenic Aβ peptides. **(B)** Schematic overview showing CM collection from APPSwe neurons treated with BACEi or a DMSO vehicle control. BACEi CM and DMSO control CM was collected from CN differentiated from Swe^WT/WT^, Swe^+/WT^, and Swe^+/+^ hiPSCs. **(C)** ELISA measurements confirm the reduction of pathogenic Aβ 1-42 production in BACEi and APP^KO^ conditions compared to the DMSO control. n=3 biological replicates per condition. **(D)** Representative immunofluorescent images of 3D microvessels after perfusion with CM collected from Swe^WT/WT^ neurons treated with DMSO (bottom left), Swe^WT/WT^ neurons treated with BACEi (bottom right), Swe^+/WT^ neurons treated with DMSO (middle left), Swe^+/WT^ neurons treated with BACEi (middle right), Swe^+/+^ neurons treated with DMSO (top left), and Swe^+/+^ neurons treated with BACEi (top right). BACEi CM results in a significant reduction in transluminal VWF fiber formation (VWF, green). Scale bars: 500 μm. **(E)** Representative IF image of 3D microvessels after perfusion with CM collected from APP^KO^ neurons. VWF transluminal fibers are absent from microvessels. Scale bars: 500 μm. **(F)** Quantification of transluminal fibers present in the microvessels. The number of transluminal fibers is significantly reduced in BACEi CM treated microvessels compared to the number of fibers formed in DMSO CM treated microvessels. n=4 biological replicates. **(G)** Association between average Aβ 1-42 peptide levels and average VWF fiber number. Dotted black line is a linear least squares regression fit to the data points that shows a strong positive linear correlation (R-squared=0.966, P value <.001). All plots: error bars, mean±SEM. *p < 0.05, **p < 0.01, and ***p < 0.001 by two-way ANOVA with Tukey’s correction for multiple comparisons test (C), linear regression (G), or one-way ANOVA with Šídák’s correction for multiple comparisons test.

Furthermore, perfusion of cerebral microvessels with Swe^+/+^ and Swe^+/WT^ BACEi CM led to the formation of only a few VWF fibers, in contrast to the significantly larger number of fibers in Swe^+/+^ and Swe^+/WT^ DMSO CM conditions (Figure 4D). Quantification of the fiber number in each media condition confirms this consistent and significant decrease across multiple replicates in BACEi treated conditions compared to the corresponding DMSO controls (Figure 4F). In our Swe^WT/WT^ control line, there was little to no fiber formation in both the BACEi and DMSO conditions as expected (Figure 4D).

To test whether VWF fibers would form in the presence of neuronal secretomes but in the complete absence of Aβ, CM was collected from APP^KO^ hiPSC-CNs (*28*). Analysis of Aβ 1-42 and 1-40 levels showed that CM from the APP^KO^ line did not secrete detectable levels of Aβ peptides as expected (Figure 4C and Supplemental Figure 4). When APP^KO^ CM was perfused through brain microvessels, no transluminal VWF fibers were observed in the vessel lumen (Figure 4E and 4F).

Notably, the number of VWF fibers in the HBMECs microvessels appears to have a positive linear correlation with the levels of Aβ 1-42, across all the media groups (Figure 4G). For example, Swe^+/+^ DMSO CM had Aβ 1-42 levels of 170 pg/mL and led to maximal fiber production of, on average, 45 VWF fibers per vessel. In contrast, Swe^+/WT^ DMSO CM had 41% less Aβ 1-42 (101 pg/mL) and 46% less average VWF fiber production (24 VWF fibers per vessel) compared to Swe^+/+^ DMSO CM, demonstrating a strong correlation between Aβ 1-42 levels and VWF fiber production (Figure 4G). Taken together, the number of fibers formed in BACEi treatment and APP^KO^ CM perfusion relative to Aβ peptide levels suggest that Aβ is a critical mediator for the formation of VWF fibers and EC activation in the engineered cerebral microvessels in response to CM perfusion.

## 3. Discussion

AD research has traditionally been focused on neuronal and glial dysfunction. Despite evidence of vascular contributions that precede the clinical diagnosis of AD, endothelial alteration in AD has been relatively poorly explored, partly due to the lack of proper humanized models. In this study, we used hiPSC-derived neurons harboring the APPSwe mutation, together with primary HBMECs and engineered 3D brain microvessels to investigate AD-associated vascular dysfunction. We validated that Swe^+/+^ mutation led to a higher concentration of Aβ deposition in CM compared to Swe^WT/WT^ control. The measured Aβ levels were in the picomolar range, which is more physiologically relevant than micromolar concentrations of exogenous Aβ peptides often applied (*16*–*18*). We demonstrated that these AD-mimicking CM caused transcriptomic and phenotypic upregulation of adhesion molecules in HBMECs. Perfusion of this high Aβ-containing Swe^+/+^ CM in 3D microvessels elicited rapid endothelial activation, following 1-hour treatment of brain microvessels, characterized by the formation of prothrombotic VWF fibers and leukocyte binding on vessel lumen. Inhibiting Aβ production in APPSwe with a BACE1 inhibitor or APP^KO^ attenuated the endothelial activation. Our findings provide direct evidence that Aβ protein in circulation induces endothelial activation and acute VWF fiber formation, which could serve as an early source of vascular dysfunction in AD.

Although it has previously been established that soluble Aβ oligomers are the primary toxic species of the three Aβ assemblies (monomers, soluble oligomers, fibrils) (*40*, *41*), our results suggest that monomeric Aβ species can drive the EC activation and dysfunction in otherwise healthy vessels. We show that the APPSwe mutation in our hiPSC-CNs led to increased Aβ 1-42 and Aβ 1-40 peptide secretion in a genotype-dependent manner (Figure 1C). However, we do not expect that Aβ oligomers or aggregates are forming in our APPSwe CM, as intracellular and extracellular Aβ aggregates were found to form only in AD-mimicking iPSC-derived neurons after long-term culture (90+ days) (*42*). This suggests that even small amounts of pathogenic Aβ peptides may lead to significant changes in vascular structure and function, which can occur early in disease progression, prior to the formation of higher-order Aβ assemblies. Our observations support increasing evidence that suggests vascular dysfunction may precede the onset of pathophysiological and cognitive symptoms in AD (*6*). Our results also support that Aβ peptides may be involved early in AD pathogenesis.

AD-related changes in brain vasculature are associated with responses to inflammatory signals, oxidative stress, adaptive immune responses, and upregulation of endothelial adhesion molecules (*43*–*47*). In particular, endothelial VCAM-1 expression (*46*) and increased binding of lymphocytes to VCAM-1 in the brains of AD patients (*48*) emphasize the role of endothelial adhesion molecules in AD. Using RNAseq, we revealed that endothelial cells respond to paracrine signals from neurons of Swe^+/+^ FAD mutation by enhancing expression of genes associated with inflammatory responses and immunoadhesion. E-selectin, ICAM-1 and VCAM-1, precursor genes for adhesion molecules in endothelial cells, were significantly upregulated in HBMECs treated with Swe^+/+^ CM. Increased expression of adhesion molecules can recruit circulating leukocytes in inflamed vessel lumens and assist infiltration of immune cells to the CNS compartment.

Facilitating neutrophil recruitment to the CNS via adhesion molecules can promote activation of microglia and astrocytes and initiate neuroinflammation and neuronal damage (*49*). CCL2 was also significantly upregulated in Swe^+/+^ CM treated HBMECs which can elicit endothelial retraction and breakdown of the blood-brain barrier, allowing for the accumulation of neurotoxic chemokines and plasma proteins (*50*). Our transcriptional analysis highlights the molecular connection between pathogenic Aβ levels, vascular inflammation, and hemodynamic dysfunction. As vascular inflammation manifests in the early stages of AD neuropathology, the inflammation cycle may exacerbate AD pathology and augment cognitive impairment.

We showed, for the first time, that perfusion of Aβ-rich CM through 3D human brain microvessels can promote acute release of VWF from endothelium and formation of ultra-large transluminal fibers in the 3D vessel lumen. We show strong linear correlation between Aβ levels in the CM and the number of transluminal VWF fibers in 3D microvessels. Perfusion of Swe^+/+^ CM with high Aβ concentrations led to large VWF fiber formation and high level of platelet binding. Attenuated Aβ 1-42 and Aβ 1-40 levels in CM with BACE1 inhibition in APPSwe CNs significantly reduced VWF fiber formation in 3D human brain microvessels. CM from APP^KO^ CNs with no detectable secretion of Aβ 1-42 and Aβ 1-40, correspondingly led to no transluminal VWF fiber formation in vessels. These data demonstrate a direct causal effect of Aβ, endothelial VWF release, and endothelial activation in human brain microvessel models.

VWF is a multimeric protein that is released from WPB either constitutively from quiescent ECs, or acutely during acute activation of ECs (*51*, *52*). Upon release from WPB, VWF unfurls its multimeric structures into elongated strings under flow, exposing crosslinking sites to bind platelets and assemble into its ultra-large fiber structure (*53*). Although how Aβ initiates WPB exocytosis is not in the scope of this work, past studies shed light on possible calcium-dependent mechanisms of Aβ induced WPB release. Transient exposure of Aβ 1-40 has been linked with disruption of calcium homeostasis in brain endothelial cells (*54*, *55*). Short-term treatment of Aβ in rat brain endothelial cells has been found to increase mitochondrial and cytosolic calcium release (*56*), which could promote exocytotic formation and exocytosis of WPBs (*57*, *58*). Release of VWF leads to fiber formation within the vessel lumens that attract platelets, bind directly to P- select glycoprotein ligan-1 on polymorphonuclear leukocytes, and recruit leukocytes for subsequent diapedesis and promote inflammatory and thrombotic events. (*59*, *60*). By comparing the perfusion outcomes of Swe^+/+^ CM with Swe^WT/WT^, Swe^+/WT^ and that with BACEi treatment and APP^KO^ media, our results demonstrated that Aβ peptides are the primary component of the CM leading to EC activation. Future human studies are needed for remaining questions such as whether there is an Aβ concentration threshold below which EC activation will not occur and above which EC dysfunction will be initiated. Understanding the minimum Aβ peptide levels required to cause EC dysfunction could help inform therapeutic strategies for AD. Overall, the results of our study have enhanced our understanding of the role of Aβ in the EC dysfunction in AD.

We have established a 3D perfusable cerebral vascular model here to investigate luminal dysfunction caused by trophic factors released from brain parenchyma in AD patients. The simplicity of only including ECs in our 3D brain vascular model allows us to identify how ECs are specifically regulated by the pathogenic Aβ released from brain parenchyma of AD patients. ECs are known to be equipped with the abluminal expression of Aβ efflux transporters such as low-density lipoprotein receptor-related protein 1 and 2 mediate elimination of Aβ from brain parenchyma to the plasma (*61*, *62*). Moreover, BBB breakdown accompanying AD progression facilitates introduction of CNS secreted pathogenic Aβ peptides on the luminal side of ECs (*63*).

There are several limitations remaining in this study. Here, we posit that BACEi inhibition and subsequent Aβ reduction leads to diminished endothelial activation. It is also believed that other cleavage products generated through the amyloidogenic cleavage of APP may be inhibited by BACE1 inhibition (secreted APPβ (sAPPβ), β carboxyterminal fragment (βCTF), and amino-terminal APP intracellular domain (AICD)). The effect of these fragments on the endothelium has not been studied (*39*, *64*, *65*) and was not examined here. This should be evaluated in future studies to better understand the effect of other APP cleavage products on endothelial cell function. Furthermore, although we have only investigated paracrine signaling from cortical neurons with APPSwe mutation, investigation of CM obtained from BBB relevant cell types like astrocytes and microglia could provide an extensive understanding of the role of other CNS cell types in AD vascular dysfunction. While we utilized the FAD APPSwe mutation, 90% of AD patients are suffering from a sporadic form of the disease and FAD is rare. Genetic studies of SAD have identified approximately two dozen associated candidate genes (*32*, *65*). Incorporating these genes into the model described here could help advance our understanding of vascular changes in sporadic AD. Finally, future studies should incorporate all relevant cell types present in the BBB. Building a comprehensive BBB model and investigating cell type specific effects on the endothelium will lead to an in-depth understanding of how cerebrovascular dysfunction occurs in AD patients.

## Methods

### Endothelial cell culture

Human brain microvascular endothelial cells (HBMECs) (CC# ACBRI 376 V; Cell systems) at passage number 3-6 were used for all experiments. HBMECs were maintained in culture flasks coated with Attachment Factor (CC# 4Z0-210; cell systems) and fed growth medium (GM) (CC# 3202; EGM-2 MV Microvascular Endothelial Cell Growth Medium-2 BulletKit (CC# 3156; EBM-2 Basal Medium), 5% fetal bovine serum (FBS), hydrocortisone, human fibroblast growth factor-beta (hFGF-B), vascular endothelial growth factor (VEGF), R3-insulin-like growth factor-1 (R3- IGF-1), ascorbic acid, human epidermal growth factor (hEGF), GA-1000 (gentamycin and amphotericin)). Cells were maintained at 37°C in a 5% CO2 incubator and media was replaced every 3 days.

### Cell lines and hiPSC neuronal differentiation

To generate AD mimicking neurons from hiPSCs we used previously established gene-edited hiPSC lines heterozygous (Swe^+/WT^) and homozygous (Swe^+/+^) for the APP Swedish mutation as well as the parental control (Swe^WT/WT^) described in Young et al. (*28*). These cell lines were generated from a previously published and characterized CV background human induced pluripotent stem cell line (*32*). This is a male cell line with an APOE ε3/ε4 genotype (*66*). A previously published APP knockout (APP^KO^) line was used as an additional control for conditioned media (CM) experiments (*28*). This cell line was made from a parental APP duplication hiPSC line and was engineered using CRISPR/Cas9 genome editing to lack APP production. Neurons from these cell lines were differentiated from hiPSCs using dual-SMAD inhibition techniques as previously described (*32*, *33*). For the experiments presented here, we started with neural precursor cells (NPCs) previously differentiated from hiPSCs and frozen. NPCs for all the APPSwe lines (Swe^+/WT^, Swe^+/+^, Swe^WT/WT^) and the APP^KO^ line were thawed and plated on a Matrigel (Growth factor reduced basement membrane matrix; CC# 356231; Corning) coated 6- well plates. NPCs were cultured in Basal Neural Maintenance Media (BNMM) + FGF (1:1 DMEM/F12 + Glutamine (CC# 11320-033; Gibco) and Neurobasal media (CC# 21103-049; Gibco), 0.5% N2 Supplement (CC# 17502-048; Gibco), 1% B27 Supplement (CC# 17504-044; Gibco), 0.5% GlutaMax (CC# 35050061; Thermo Fisher Scientific), 0.5% insulin-transferrin-selenium (CC# 41400045; Thermo Fisher Scientific), 0.5% NEAA (CC# 11140050; Thermo Fisher Scientific), 0.2% b-mercaptoethanol (CC# 21985023, Life Technologies) + 20 ng/mL FGF (R&D Systems, Minneapolis, MN)). The NPCs were allowed to reach 70% confluence for 1-3 days with media changes every other day. Once the NPCs were sufficiently confluent, cells were dissociated with Accutase (CC# 07930, STEMCELL Technologies) and passaged into Matrigel- coated 10 cm culture plates for neural differentiation. 24 hours after replating, the cells were switched to Neural Differentiation media (BNMM + 0.2 mg/mL brain-derived neurotrophic factor (CC# 450-02; PeproTech) + 0.2 mg/mL glial-cell-derived neurotrophic factor (CC# 450-10; PeproTech) + 0.5 M dbcAMP (CC# D0260; Sigma Aldrich). Media was changed twice a week for 21 days at which point the differentiation is considered finished. Cells were maintained at 37°C in a 5% CO2 incubator.

### Conditioned media preparation

Following 21 days of differentiation, Neural Differentiation media was refreshed and was then kept on the cells for at least 72 hours with no media change to generate conditioned media (CM) for 2D and 3D treatment experiments. After 72 hours, the media was collected and stored at 4°C for short-term storage and −20°C for long-term storage. To generate CM from BACEi treated APPSwe neurons, neurons were dissociated from 10 cm plates using Accutase following 21 days of differentiation and split into 3 wells of a Matrigel-coated 6 well plate. This was repeated for all cell lines. At least 3 days later, neurons were treated with either 25 nM β-secretase inhibitor (CC# HY- 13240; BACEi LY2886721; MedChemExpress) or dimethyl sulfoxide (DMSO, as vehicle control) or left untreated every 24 hours 72 hours with no media change. After 72 hours of treatment, media from untreated, DMSO, and BACEi treated neurons was harvested for perfusion through microvessels, treatment in 2D, and quantification of Aβ1–40 and Aβ1–42 peptides secreted by neurons.

### Amyloid Beta Measurements

Aβ1–40 and Aβ1–42 peptides were measured using the methods previously described by Young et al., 2015. (*34*) Conditioned media collected from the Swe^+/WT^, Swe^+/+^, Swe^WT/WT^, and APP^KO^ cell lines after a 3-week differentiation were frozen at −20°C until use. The media was run on an Aβ Triplex ELISA plate (CC# 151200E-2; Meso Scale Discovery).

### 3D cerebral microvessel fabrication

Engineered microvessels were fabricated using soft lithography and injection molding techniques previously described by Zheng et al., 2012. Briefly, type I rat tail collagen was dissolved in 0.1% acetic acid at a stock concentration of 15 mg/mL (*36*). The collagen was neutralized and diluted to 7.5 mg/mL on ice with 1 M NaOH (20 mM final), 10x M199 (CC# 11825015; Thermo Fisher Scientific), and GM. A microfabricated polydimethylsiloxane (PDMS) stamp was used to define a double-grid vessel network with a feature height of 150 μm. Collagen was molded around this PDMS stamp. A flat collagen gel was also generated. After gelation at 37°C, the channels were incubated with the flat collagen gel to generate closed off microchannels that can be perfused and seeded with endothelial cells (ECs). These two gels merged together through additive bonding upon incubation at 37°C. Once the gel was fabricated, HBMECs were lifted from culture flasks with trypsin (CC# MT-25-052-CI; Thermo Fisher Scientific) and resuspended in GM at a concentration of 7-10×10^6^ cells/mL. Using a gel loading pipette tip, 10 uL of this cell suspension is injected into the inlet of the microvessel and allowing them to circumferentially attach to the collagen gel under static conditions for 1 hour prior to the start of gravity-driven flow. Microvessel constructs were cultured under gravity-driven flow for 5-7 days with media replenishment approximately every 12 hours.

### Conditioned media treatment

For the treatment of HBMECs with CM in 2D, HBMECs were plated at a density of 0.03M/well in a glass bottom chamber slide (CC# 80827, Ibidi) and allowed to recover for 48 hours prior to treatment. Prior to treatment, the cells were washed with EBM-2 basal media (serum-free) for 5 mins. Then, 250 uL of CM per well was then added to the cells for 8 hours, following which the cells were fixed in 4% PFA. To treat our engineered microvessels in 3D, vessel fabrication was performed as described above. Following vessel fabrication and culture for 5-7 days under gravity-driven flow, the vessels were treated with CM from all of our APPSwe lines and CM from BACEi treated neurons. Vessels were washed with EBM-2 basal media (serum-free) for 20 mins. Vessels were then perfused with CM for 1 hour. Media was replenished every 20 mins to continue gravity-driven flow throughout the duration of treatment. Following 1 hour of perfusion, the vessels were fixed with 4%PFA for 20 mins for downstream analysis and washed with phosphate buffered saline (PBS) three times for 20 mins each. For RNAseq analysis, vessels were treated for 8 hours with media replenished every 2 hours.

### RNA isolation and RNAseq analysis

RNA was collected from two different culture subtypes (2D cultures and 3D microvessels) for three different media conditions (unconditioned BNMN media, conditioned control media and conditioned Swe^+/+^ media). RNA from 2D cultures was collected following media treatment from 24 well plates by addition of RLT lysis buffer and pipette homogenization. RNA from 3D vessels was collected by perfusion of RLT lysis buffer in the inlets and collection of the lysates in the outlets. Lysates from three wells in 2D cultures were pooled as one sample replicate, and two vessels from 3D cultures were pooled as one replicate. Total RNA was isolated from the lysate using on-column digestion of genomic DNA via RNeasy Micro Kit (CC# 74004; Qiagen). RNA quality was assessed using High sensitivity RNA TapeStation 4200 (Agilent Technologies) and total RNA samples with RNA integrity number >8 were used to proceed to library preparation. For sequencing, cDNA preparation was performed using Takara SMART-Seq v4 Ultra Low Input RNA Kit (SMARTv4), library preparation using Illumina Nextera XT DNA Library Prep Kit (Nextera XT) and sequenced with Illumina NovaSeq 6000 (paired-end 50 cycles). Sequenced reads were aligned using STAR2 aligner and gene-level raw counts was obtained using STAR2’s internal quantification method. Gene ontology analysis was performed using iDEP.951 for pathway analysis of differentially expressed genes.

### Isolated platelet and peripheral blood mononuclear cell (PBMC) perfusion in microvessels

Whole blood was drawn from normal healthy donors into a blood collection tube (BD, CC# 364606; Vacutainer) containing 1.5mL of acid citrate dextrose (ACD). Platelets were isolated from whole blood using an existing protocol provided by the manufacturer. Briefly, collected blood was centrifuged at 200g for 20 minutes (acceleration 4; deceleration 0) at room temperature for platelet-rich plasma (PRP) extraction. PRP was subsequently centrifuged at 1200g for 10 minutes to obtain pellets containing the isolated platelets. Pellets were resuspended in CGS buffer containing 120mM NaCl, 13mM Sodium Citrate and 30mM Glucose supplemented with Prostaglandin E1 (sigma P5515-1MG) at 1:10,000 (vol: vol) to inhibit aggregation of platelets. Isolated platelets rinsed in CGS/PGE1 buffer were counted under a bright-field microscope using a hemocytometer and centrifuged again using the previous setting. Pellets were resuspended carefully in Tyrode’s buffer containing 5.5mM glucose, 10mM HEPES, 138mM NaCl, 12mm HaHCO3, 2.9 mM KCL and 0.36mM NaH2PO4 and stained with CD41a (CC# 555467; BD) conjugated with phycoerythrin (PE) for 30 minutes. Following staining, platelet concentration was adjusted to 5 x 10^7 platelets/mL by supplementing an additional Tyrode’s buffer.

Isolated platelets were perfused in conditioned media treated microvessels to assess the functional activation of endothelial cells. CD41a labeled isolated platelets were perfused in HBMEC microvessels treated with conditioned media for 1 hour through transfer pipettes to avoid shear-induced activation. Approximately 150 μl of platelets resuspended in Tyrode’s buffer was pipetted into the inlets of microvessels and incubated at 37C for 20 minutes. Following platelet treatment, vessels were washed three times, each rinse for five minutes, to eliminate residual unbound platelets within the vessel lumen. Immediately after the wash step, microvessels were fixed in 4% paraformaldehyde solution and rinsed with PBS three times. Platelet adhesion analysis was performed after performing immunofluorescent staining of VWF and confocal imaging.

PBMCs were also isolated from freshly drawn blood samples collected in ACD blood collection tube (BD CC# 364606; Vacutainer). PBMCs were isolated using density gradient centrifugation in a 50mL SepMate tube (StemCell Technologies) following manufacturer’s protocol. PBMCs were stained with pan-leukocyte marker CD45 (CC# 555485; BD biosciences) for live imaging of leukocyte adhesion on Swe^+/+^ CM treated HBMEC microvessel for 1 hour.

### Scanning electron microscopy imaging of platelet bound VWF fibers

Swe^+/+^ perfused microvessels were fixed in ½ strength Karnovsky’s fixative (2% paraformaldehyde and 2.5% glutaraldehyde) overnight at 4C. The top and bottom layers of the collagen matrices were gently teased apart and rinsed with 0.1M cacodylate buffer. Samples were dehydrated with graded series of alcohols and critical point dried (Autosamdri, Tousimis Corp). Microvessel layers were sputter coated with Au/Pd alloy (Denton Desk IV, Denton Vacuum) and imaged using a JSM 6610 LV scanning electron microscope (JEOL) at an accelerating voltage of 5kV.

### Flow cytometry analysis

To characterize vascular cell adhesion molecule 1 expression on the surface of endothelial cell following CM treatment, multiparameter FACS analysis of CM treated human brain endothelial cells was performed. BD FACS Canto II was used to analyze VCAM-1 positive cells using FTIC- anti-VCAM-1 antibody (CC# ab24853; Abcam) and the number of positive cells were quantified using with FlowJo software (Tree Star). Unstained controls were used to compensate and adjust voltage.

### Immunofluorescent staining and image analysis

At the end of each experimental time point, both microvessels and 2D endothelial cultures were fixed with 4% paraformaldehyde for 20 minutes and washed with PBS three times. For immunofluorescent staining of microvessels, reagents were perfused through the microchannels through the inlet of the acrylic jigs. PBS containing 2% bovine serum albumin (BSA) was used to block microvessels and 2D cultures for analysis of extracellular VWF. Following 1 hour of blocking, FITC conjugated VWF antibody (1:100; Abcam CC# ab8822) was added to samples for 1 hour at room temperature and washed three times with PBS before proceeding to stain with the remaining antibody of interest. Treatment with 2% BSA with 0.1% Triton-x preceded before staining with antibodies including CD41a (CC# 555467; BD), phalloidin (CC# A12380; Thermo Fisher), CD144 (CC# 136008; BioLegend) for cell permeabilization and blocking non-specific binding. All stained microvessels and 2D cultures were imaged using Yokogawa W1 spinning disk confocal microscope or Leica SP8 laser confocal microscope. For microvessels, the entire collagen construct was imaged using a 5 by 5 large z stack image stitch (10% blending stitch) under a 10X objective with a z-step size of 5.5 μm. 3D reconstruction and cross-sectional images of the microvessels were rendered using ImageJ software.

### VWF fiber quantification

VWF fibers were quantified into two distinct wall-fiber and transluminal fiber categories. Transluminal fibers were quantified from fibers formed cross-sectionally within the microchannels and are characteristically thicker and straighter than wall fibers. Wall fibers were quantified from string fibers found circumferentially on the endothelial surface and are smaller and tortuous by nature. For classification of the number of fibers found in each group, stitched z stack Images of each vessel were divided into three layers: top, center lumen and bottom. The top and bottom layers were constructed from compilation of initial and final 4 z-stacks (~20μm) of each image construct respectively. The luminal layer was constructed from compiled z stack images excluding the initial and final 4 layers.

### Statistical analysis

Statistical analysis of fiber quantification was performed with either one-way ANOVA or two-way ANOVA for comparison between groups followed by Tukey’s or Šídák’s post hoc test for multiple comparisons between groups. Statistical analysis on the RNAseq was conducted with Fischer’s Exact test. Significance was defined as a value of p < 0.05. For all imaging experiments the data was analyzed in a blinded manner. Statistical analysis was performed using GraphPad Prism software. Information on statistical details of individual experiments can be found in the respective figure legends.

## Acknowledgments

We would like to acknowledge the Lynn and Mike Garvey Imaging Laboratory in the Institute for Stem Cells and Regenerative Medicine (ISCRM) and the Flow Cytometry Core at the University of Washington. We also acknowledge the Electron Microscopy Laboratory and Genomics & Bioinformatics Laboratory both at Fred Hutchinson Cancer Research Center. We would also like to acknowledge that schematics were made using BioRender.

## Funding

We acknowledge the financial support of National Institute of Health grants R21AG074373 (to Y.Z. and J.E.Y), R01AG062148 (to J.E.Y) and T32AG066574 (to K.M.E, training grant). Additionally, we acknowledge the financial support of ISCRM (to Y.S., fellowship).

## Author contributions

Conceptualization, Y.Z., J.E.Y., Y.S., and K.M.E.; Methodology, Y.Z., J.E.Y., Y.S., K.M.E., and C.H.; Investigation, Y.Z., J.E.Y., Y.S., K.M.E, C.H., S.J., and M.S.M.; Writing – Original Draft, Y.Z., J.E.Y., Y.S., and K.M.E.; Writing –Review & Editing, Y.Z., J.E.Y., Y.S., K.M.E., C.H., J.X., S.J., and M.S.M.; Funding Acquisition, Y.Z, J.E.Y., Y.S., and K.E.; Supervision, Y.Z. and J.E.Y.

## Competing interests

The authors declare that they have no competing interest.

## Data and materials availability

All data needed to evaluate the conclusions in this paper are present in the main paper and/or the Supplementary Materials. Additional data related to this paper may be requested from the corresponding author.

## Notes

### Competing Interest Statement

The authors have declared no competing interest.

