## Supplemental Materials for "Amyloid beta peptides (Aβ) from Alzheimer’s disease neuronal secretome induce endothelial activation in a human cerebral microvessel model"

### Supplemental Figure 1.

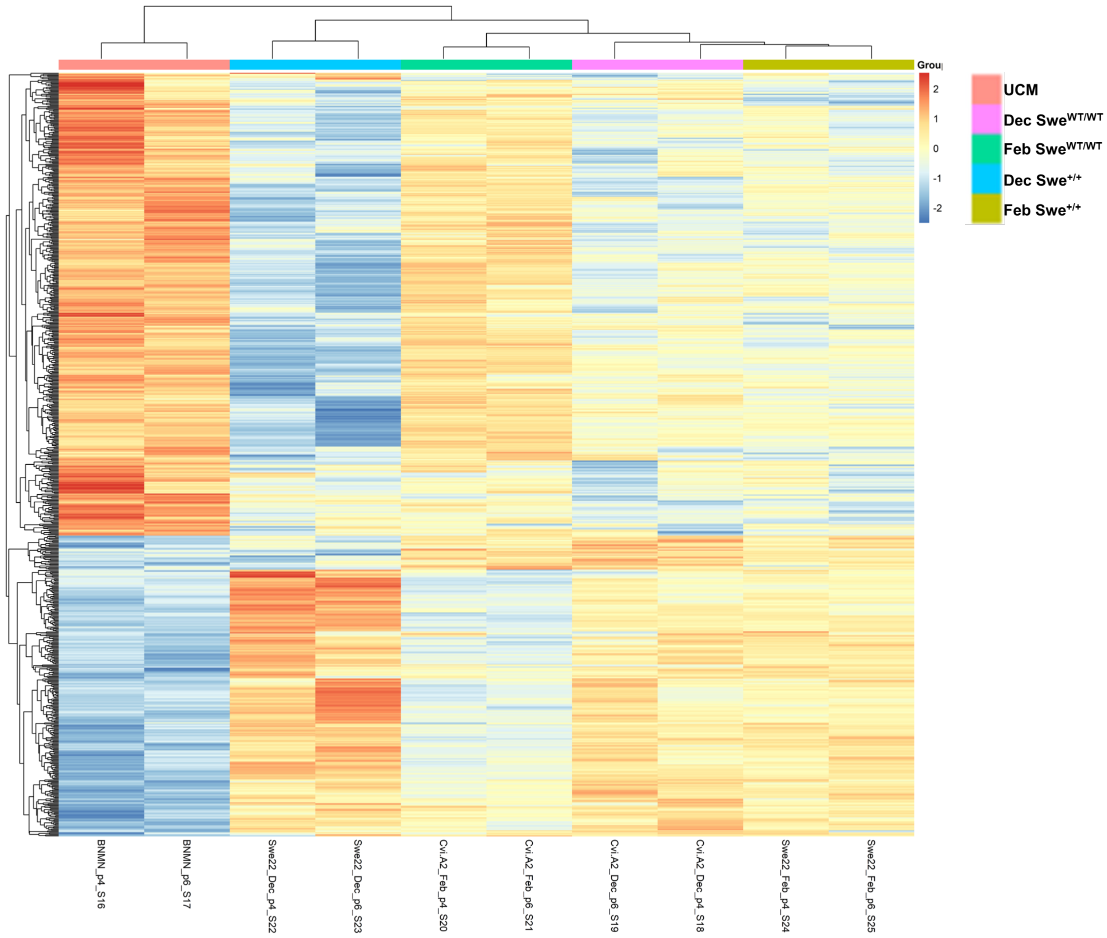


**Figure S1. Bulk RNAseq comparison showing gene expression profile in conditioned media treated HBMECs.** Heatmap of log counts per million (CPM) values of differentially expressed genes in HBMECs treated with UCM, Swe$^{WT/WT}$, and Swe$^{+/+}$ CMs.

### Supplemental Figure 2.

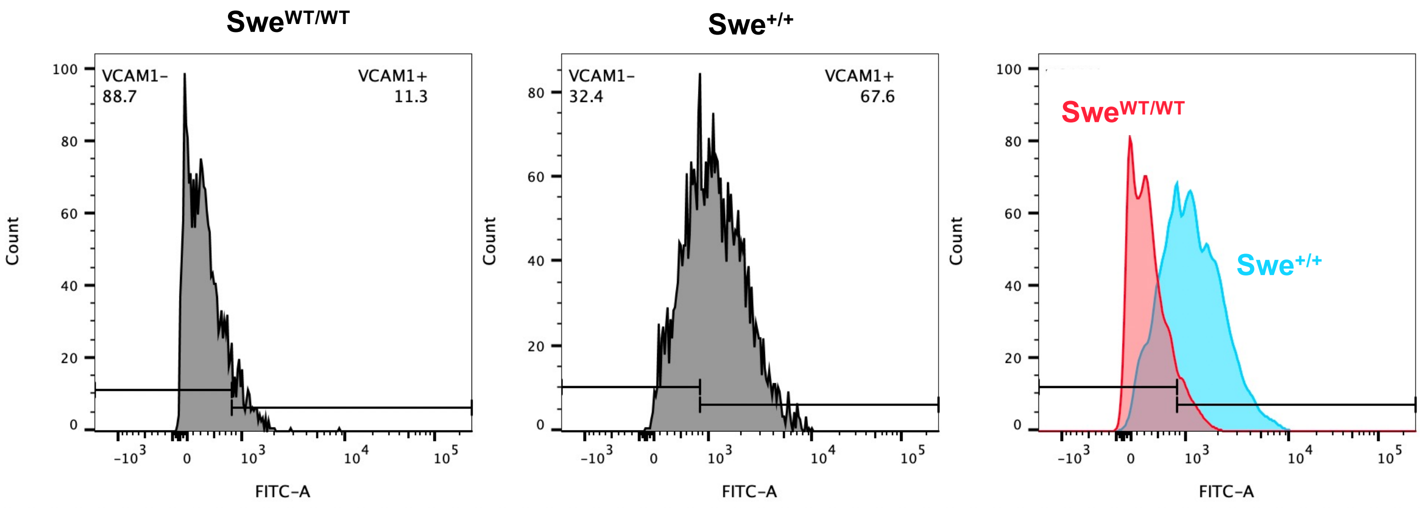


**Figure S2. Flow cytometry analysis of VCAM-1 (FITC) expression on HBMECs.** 2D HBMECs treated with $Swe^{WT/WT}$ CM and $Swe^{+/+}$ CM at 37 °C for 24 hours were stained with VCAM-1 analyzed with flow cytometry. HBMECs treated with $Swe^{+/+}$ CM results in enhanced expression of VCAM-1.

**Supplemental Figure 3.**

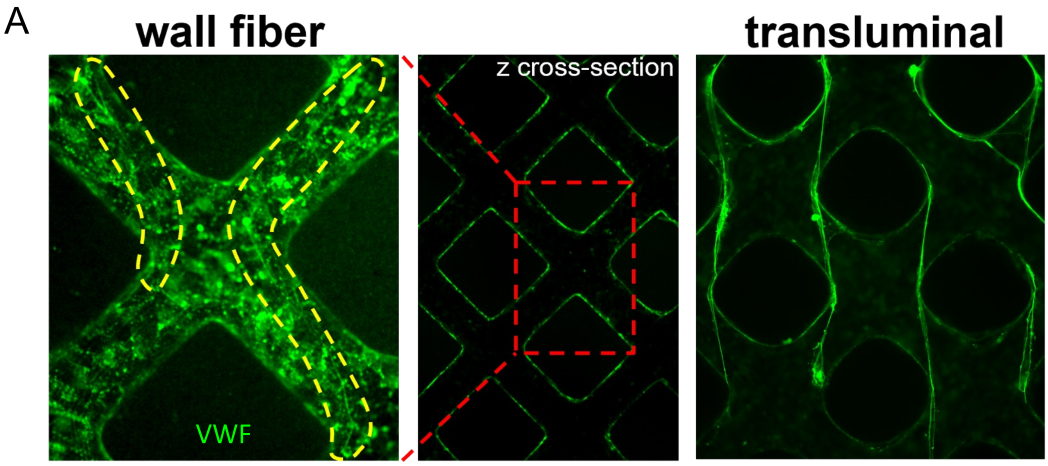


**Figure S3. Identification of wall and transluminal fibers following CM perfusion in microvessels.** Wall fibers are classified as fibers found on the surface of endothelial cells that line the microvessels. Transluminal fibers are classified as thick fibers present across the lumen of the microvessels.

**Supplemental Figure 4**.

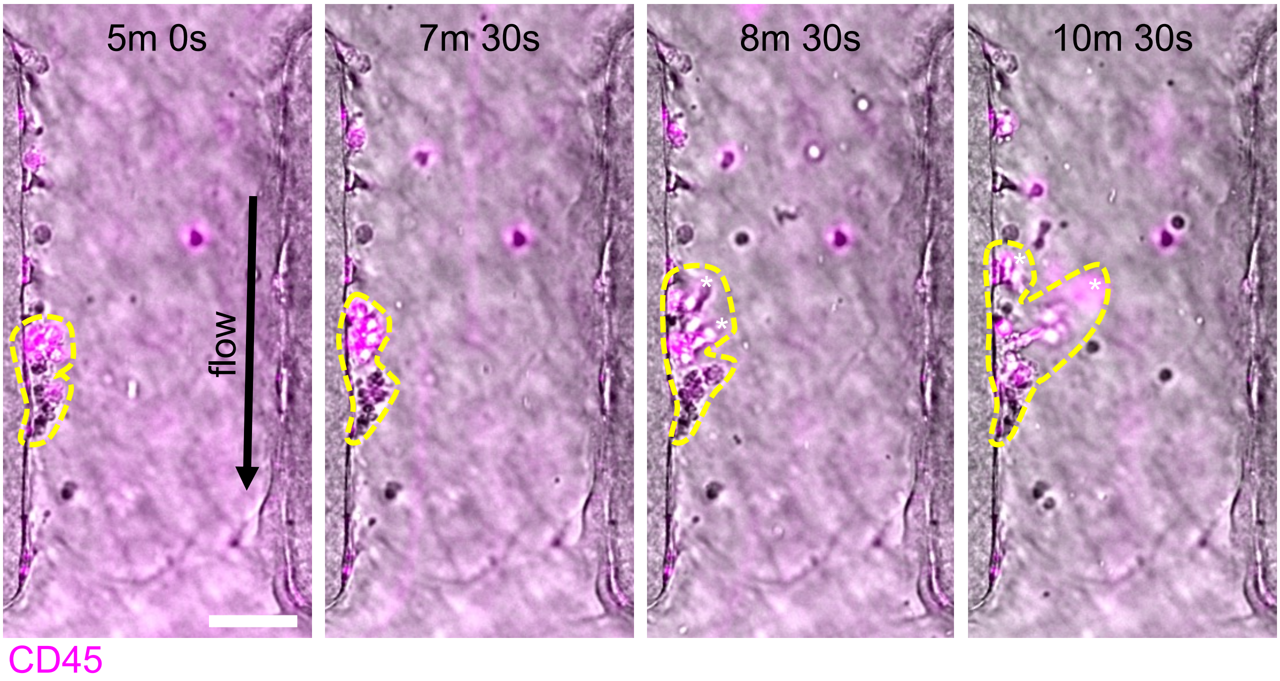


**Figure S4. Leukocyte tethering and rolling in Swe$^{+/+}$ CM treated microvessels**. Time-lapse images show a cluster of isolated leukocytes bound to activated HBMEC microvessels under flow (black arrow). The leukocytes tether and roll (yellow dashed outline) on activated ECs against the direction of flow. Scale bar: 50 µm.

**Supplemental Figure 5.**

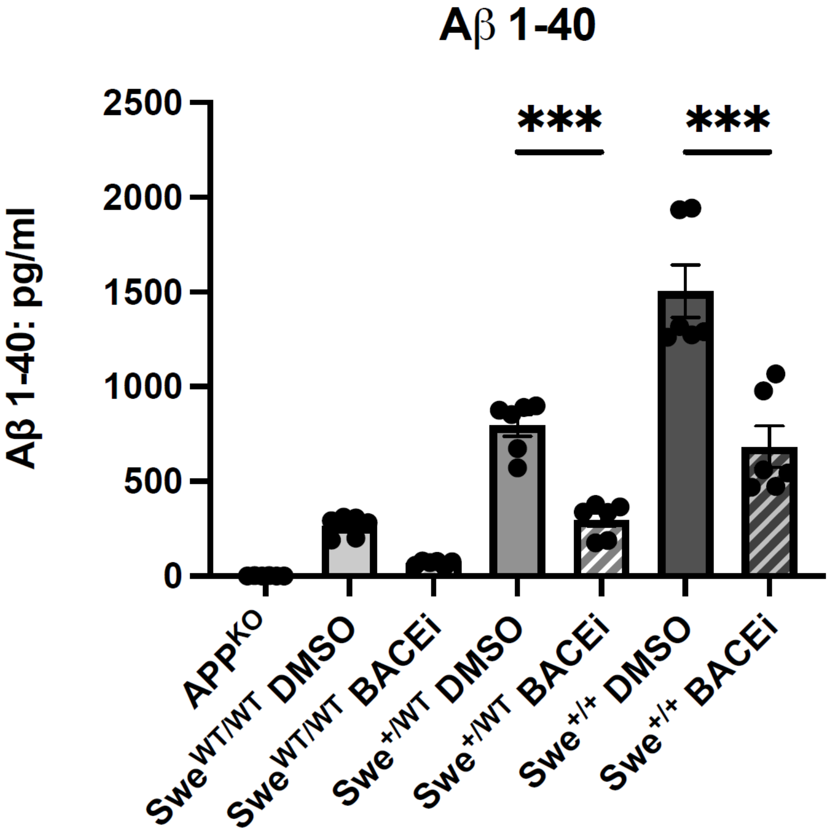


**Figure S5. ELISA measurements confirm the reduction of Aβ 1-40 production in BACEi and APP$^{KO}$ conditions compared to the DMSO control.** n=3 biological replicates per condition. Error bars, mean ± SEM. *$p < 0.05$, **$p < 0.01$, and ***$p < 0.001$ by two-way ANOVA with Tukey's correction for multiple comparisons test.
